# The central clock suffices to drive the majority of circulatory metabolic rhythms

**DOI:** 10.1101/2022.01.24.477514

**Authors:** Paul Petrus, Jacob G. Smith, Kevin B. Koronowski, Siwei Chen, Tomoki Sato, Carolina M. Greco, Thomas Mortimer, Patrick-Simon Welz, Valentina Zinna, Kohei Shimaji, Marlene Cervantes, Pierre Baldi, Pura Muñoz-Cánoves, Paolo Sassone-Corsi, Salvador Aznar Benitah

**Author notes:** Authors contributed equally (Pura Muñoz-Cánoves, Paolo Sassone-Corsi, Salvador Aznar Benitah).

## Abstract

Life on Earth anticipates recurring 24-h environmental cycles via genetically-encoded molecular clocks active in all mammalian organs. Communication between these clocks is believed to control circadian homeostasis. Metabolism can be considered a form of inter-tissue communication language that results in temporal coordination of systemic metabolism between tissues. Here we characterize the extent to which clocks in different organs employ this means of communication, an area which remains largely unexplored. For this, we analysed the metabolome of serum from mice with tissue-specific expression of the clock gene *Bmal1*. Notably, having functional hepatic and muscle clocks can only drive a minority (13%) of the oscillating metabolites in circulation. Conversely, limiting *Bmal1* expression to *Syt10*- expressing neurons (which are enriched in the suprachiasmatic nucleus [SCN], the master pacemaker that regulates circadian rhythms) restores rhythms to 57% of circulatory metabolites and 28% of liver transcripts, and rescues glucose intolerance. Importantly, these parameters were also restored in clock-less mice upon rhythmic feeding, indicating that the central clock mainly regulates metabolic rhythms via behavior. These findings explicate the circadian communication between tissues and highlight the importance of the central clock in governing those signals.

## Main

Throughout evolution the Earth has spun around its own axis with a 24-h cycle. Thus, life has adapted to the re-occurring environmental fluctuations that accompany Earth’s rotation with a genetically encoded molecular clock, henceforth: the core clock machinery (*1*). These clocks exist in every organ in the mammalian body and are believed to cooperate in order to control circadian homeostasis (*2*). A way of inter-organ communication is via metabolic fluctuations which is believed to function similar to an economic system taking the supply and demand of various tissues into consideration (*3, 4*). In fact, systemic metabolic homeostasis is temporally coordinated between tissues and is perturbed when challenged with a high fat diet (*5*). Nevertheless, the contribution of specific tissue clocks in regulating systemic metabolic rhythms remains a virtually unexplored field.

To address the degree to which specific-tissue clocks can regulate systemic metabolism, we first analyzed the role of local clocks in driving the strong temporal metabolic coherence that exists between serum, liver and muscle (*5*). Metabolic rhythms are intimately linked with the core clock machinery, which comprises of a transcriptional–translational feedback loop: the transcription factors BMAL1 and CLOCK heterodimerize and bind to E-box elements, thereby transcribing an array of genes that includes their own repressors *Period1-3* (*Per1-3*) *and Cryptochorme1-2* (*Cry1- 2*) (*6, 7*). This feedback loop takes about 24 h to complete and integrates metabolic input and output, to control circadian homeostasis (*8*). We used our previously-generated conditional gene trap approach (*9, 10*), in which tissue-specific expression of *Bmal1* is achieved through Cre-mediated excision of a stop-FL cassette between exon 5 and 6 of *Bmal1* in mice. Expression of Cre recombinase under the *Alfp*- or *Hsa*- promotor results in selective *Bmal1* expression in liver (*9*) (liver-RE) or skeletal muscle (muscle-RE), respectively (see accompanied manuscript Kumar et.al.). Furthermore, mice without *Bmal1*-stop-FL are wild-type (WT) for *Bmal1* (e.g., it is expressed in all tissues), while Cre-negative *Bmal1*-stop-FL mice are knockout (KO) for *Bmal1* organism-wide. Mice were sacrificed every 4 h during a 24-h diurnal cycle and serum was collected from all genotypes. The serum samples were analyzed by global metabolomics using liquid chromatography–mass spectrometry (LC/MS).

In WT mice, 25% (217/868) of detected circulating metabolites displayed significant circadian oscillations (Table S1, JTK cycle (*11*); *P* < 0.05). *Bmal1*-KO mice lost rhythmicity of all metabolites observed for WT except for cysteine-S-sulfate (Fig. 1A), confirming that *Bmal1* expression is crucial for oscillations in circulatory metabolites. Reconstitution of the hepatic clock (liver-RE) or muscle clock (muscle-RE mice) restored a unique set of metabolites by each tissue; strikingly, however, these metabolites only comprised ∼13% of the WT oscillating metabolome (Fig. 1B). These data suggest that local peripheral clocks in isolation are insufficient to drive the majority of the circadian metabolic output into circulation. Of note, similar to *Bmal1*-KO mice, liver-RE and muscle-RE mice lacked detectable feeding–fasting cycles, and daily differences in locomotor activity were dramatically reduced as compared to WT (*9*) (see accompanied manuscript Kumar et.al.). Nevertheless, the oscillating metabolites common between WT and liver-RE or muscle-RE mice displayed a similar circadian pattern as in WT mice (Fig. 1B), with aligned phases (Fig. 1D) and amplitudes (Fig. 1E). These metabolites comprised intermediates in the histidine-, nicotinamide- and topical agent pathways for the liver clock, and polyunsaturated acyl carnitine for the muscle clock (Fig. 1F). Of note, regulation of nicotinamide metabolism provides a link to the clock system: expression of *Nampt* is controlled via the core clock machinery and generates nicotinamide adenine dinucleotide (NAD^+^) through the salvage pathway (*12*), which feedback to regulate the clock machinery via SIRT1 activity (*13*). In addition, histidine metabolism has been linked to circadian rhythms in the brain (*14*), presenting a potential role of the hepatic clock in regulating central functions. However, the autonomous liver clock is unlikely to regulate behavioral rhythms, as liver-RE mice display arrhythmic behaviors of feeding and locomotor activity (*9*). Furthermore, the muscle clock is necessary for local lipid metabolism, including for long-chain polyunsaturated fatty acid metabolism (*15*); data from the muscle-RE serum demonstrated partial sufficiency of this metabolic pathway. In sum, autonomous liver and muscle clocks in isolation are sufficient to drive only a small number of metabolic rhythms in circulation, revealing that the majority of circulatory metabolites are dependent on other tissue clocks, or communication between tissue clocks.

**Figure 1.**
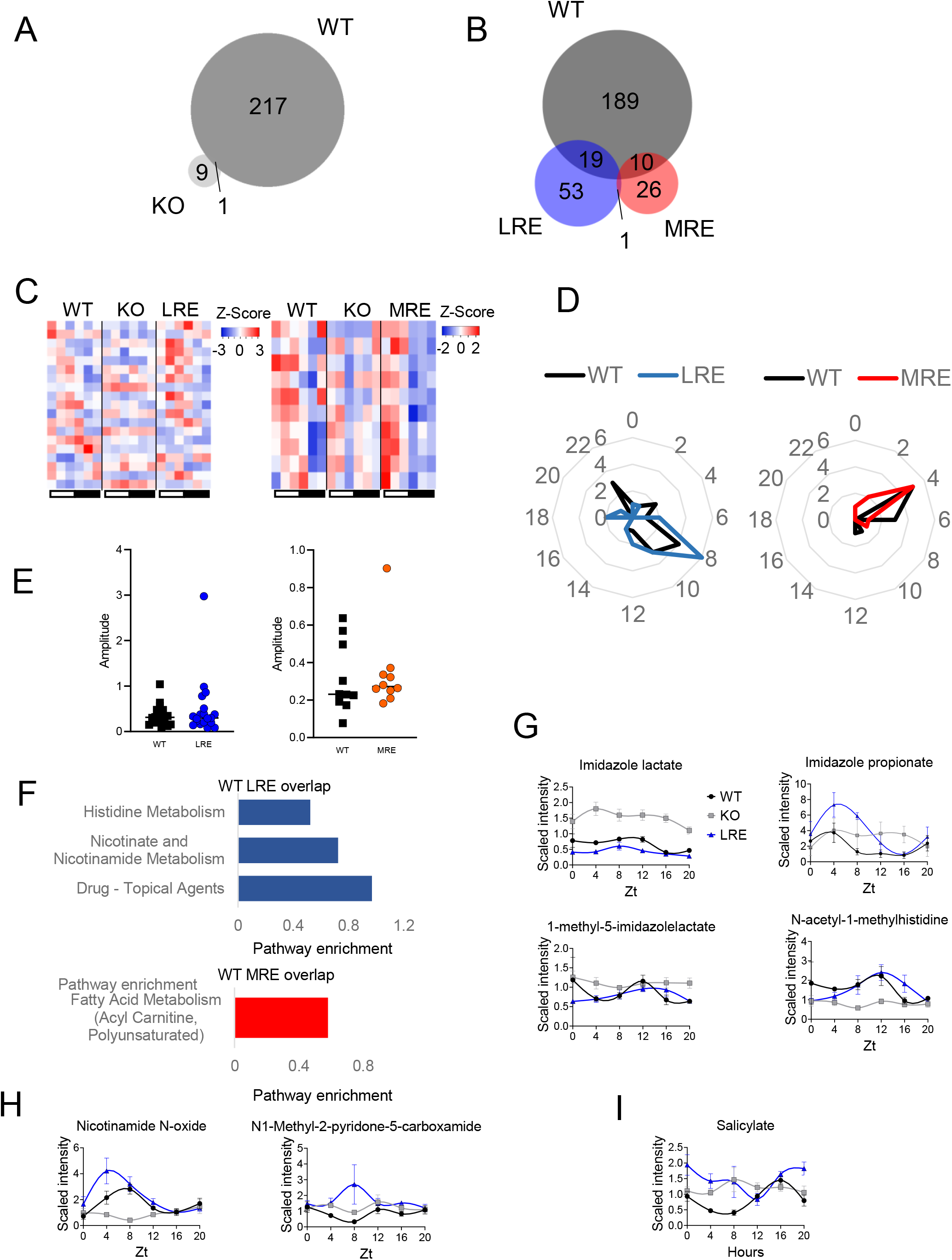

Metabolic rhythms require substrates via food intake or from stores within the body (*4*). Thus, rhythmic feeding behavior is likely a key determinant of systemic metabolic oscillations. Behavioral rhythms are controlled by the central clock of the SCN (*16*). The *Syt10* gene is highly expressed in the central circadian pacemaker located within the SCN, along with other brain regions (Extended Data Fig. 1A) (*17, 18*). Importantly, knocking out *Bmal1* in *Syt10*-expressing neurons results in arrhythmic behavior when the mice are placed in constant darkness (*18*). To study the degree to which centrally-expressed *Bmal1* is sufficient for generating systemic metabolic rhythms, we crossed the stop-FL mice with mice expressing the Cre recombinase under control of the *Syt10* promotor (hereafter referred to as brain-RE). Indeed, BMAL1 was successfully expressed in the brains of brain-RE mice, including within the SCN, but also in other brain regions such as the prefrontal cortex and hippocampus. Importantly, BMAL1 was not expressed in peripheral tissues of the brain-RE mice (Extended Data Fig. 1B–F).

To assess if BMAL1 expression in Syt10 neurons restored behavioral rhythms, we next analyzed locomotor activity, metabolism and feeding rhythms for brain-RE mice as compared to WT or KO mice. As expected, circadian rhythms of behavior and metabolism were partially restored in the brain-RE mice and maintained rhythmicity in constant darkness (Extended Data Fig. 2A-C). Partial restoration implies that clocks in other cell types of the brain, and potentially peripheral clocks as well, are required for full robustness of behavioral rhythms (see accompanied manuscript Kumar et.al.). Despite partially rescued behavior, brain-RE mice displayed a remarkable number of metabolite oscillations in serum, with a 57% overlap with those in WT (e.g., of the 223 metabolites in WT mice, 124 were found in brain-RE) (Fig. 2A), suggesting that the central clock governs processes which define the majority of systemic metabolic rhythms. Oscillating metabolites common to WT and BRE mice exhibited similar phases; however, most daytime metabolites peaked with a 4-h delay in the brain-RE as compared to WT mice (Fig. 2B, C). Strikingly, in brain-RE mice, 174 (of 298) metabolites gained *de novo* oscillation (Fig. 2A), and the amplitudes of commonly oscillating metabolites were significantly reduced, as compared to WT mice (Fig. 2D). These results suggest that peripheral clocks (and/or other brain clocks) are able to suppress aberrantly oscillating metabolites, whilst also modulating the amplitudes and phase of others.

**Figure 2.**
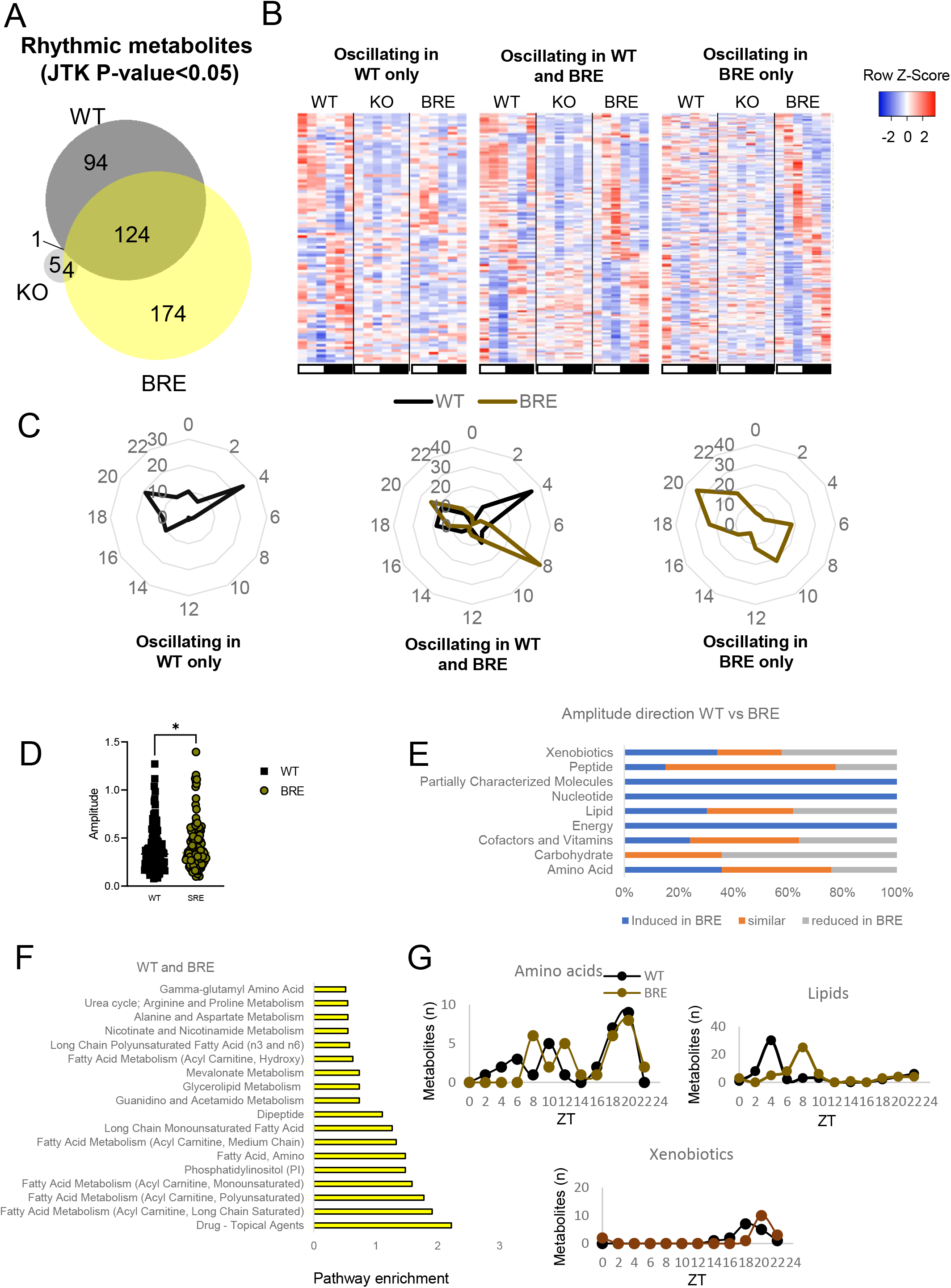

The metabolic classes that displayed increased amplitudes in brain-RE mice included nucleotide- and energy-related metabolites, as well as partially characterized molecules (Fig. 2E). In contrast, carbohydrates displayed mainly a reduced amplitude in brain-RE mice (Fig. 2E). Although the hepatic and muscle clocks alone cannot drive systemic oscillations of carbohydrate metabolites, they are necessary to buffer these rhythms (*19, 20*). The commonly oscillating metabolites were enriched for various acyl carnitines and the urea cycle, along with metabolites in the drug–topical agent class (Fig. 2F and Extended Data Fig. 3). Ureagenesis is controlled by the core clock protein CLOCK (*21*), suggesting that components of the clock machinery may continue to control circadian metabolic homeostasis possibly via other unknown feedback loops. The uniquely oscillating metabolites in WT or brain-RE mice were enriched for several fatty acid metabolic classes or sphingomyelin metabolism, respectively (Extended Data Fig. 4A, B). Most oscillating metabolites belonged to the amino acid, lipid and xenobiotic metabolic classes; oscillation patterns of amino acids and xenobiotics were similar between WT and brain-RE, while that of lipids peaked with a 4-h delay in brain-RE mice (Fig. 2G). Taken together, these data suggest that the central clock is sufficient to drive most circadian circulating metabolic rhythms, yet phase and amplitude often requiring additional regulation by other clocks.

We next investigated to what degree the central clock suffices in regulating transcriptional oscillations of the periphery in the absence of peripheral clocks. We thus studied the hepatic rhythmic transcriptome from brain-RE mice by performing RNA-seq over a 24-h period (Table S2). Of the 3392 oscillating genes in WT livers, 923 oscillated in brain-RE livers (Fig. 3A). Thus, the central clock is sufficient for driving ∼27% of the hepatic transcriptome, which is higher than the ∼13% driven by the local autonomous liver clock (*9*). Interestingly, the liver-RE mice displayed a higher recovery (∼41%) of the circadian liver transcriptome once feeding rhythms were introduced (*22*), suggesting that regulation by both the local clock and by feeding rhythms were additive rather than redundant (27%+13%=41%). The transcription phases were distributed throughout the day in WT livers, while most hepatic mRNA peak at ZT6 and ZT18 in brain-RE mice (Fig. 3B, C), possibly in direct response to the feed/fasting cycles. In line with the serum metabolite rhythms, the amplitude of hepatic transcripts was higher in brain-RE than WT mice (Fig. 3D). The commonly oscillating transcripts were enriched for genes involved in the peroxisome pathway and the circadian clock system (Fig. 3E, Extended Data Fig. 5A, B). The former is in agreement with the oscillations observed in the acyl carnitine pathway (Fig. 2F), suggesting that fatty acid beta-oxidation in peroxisomes, which results in excessive acyl-carnitine, is under central circadian control. Of note, oscillations of the circadian genes *Per1* and *Per2* were similar in WT and brain-RE livers, suggesting that their transcriptional control can be uncoupled from hepatic *Bmal1* (Extended Data Fig. 5A) and is likely driven by systemic signals (*23*). In addition, 2,425 genes were only rhythmic in WT mice (Fig. 3A), including ones involved in VEGFA signaling and HIF-1-alpha, in the IFN-gamma pathway and the methionine cycle (Extended Data Fig. 6A, Extended Data Fig. 7A, B). These pathways have all been linked to the clock system (*24*–*26*) and require peripheral clocks to maintain circadian homeostasis. *De novo* oscillating transcripts in the brain-RE livers were strongly enriched for genes in the branched- chain amino acid pathway (Extended Data Fig. 6B, Extended Data Fig. 8); however, oscillations in circulating metabolites in this pathway were not enriched in brain-RE mice (Fig. 2F).

**Figure 3.**
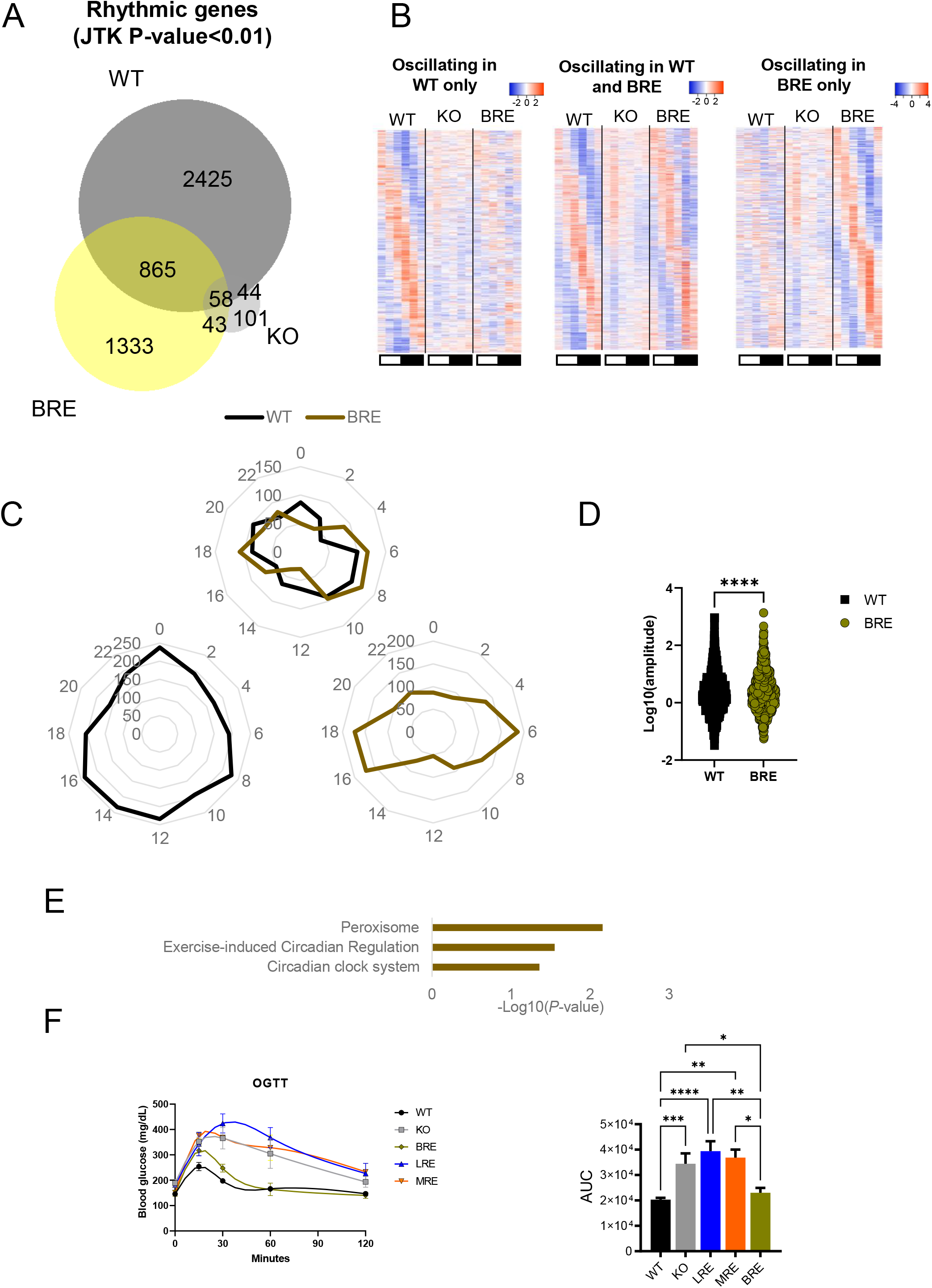

Glucose metabolism is controlled by the circadian system, and systemic glucose homeostasis requires clocks in multiple organs (*4*), but the degree to which each clock suffices is unclear. In fact, we observed a rescue of acyl carnitine oscillations in brain-RE mice which may potentially play a role in linking fatty acid oxidation and glucose homeostasis (*27*). Hence, we hypothesized that the brain clock regulates glucose homeostasis independent of peripheral clocks. Thus, we performed oral glucose tolerance tests in all RE genotypes presented herein. Bmal1-KO mice displayed a poor glucose tolerance, and this was not improved by reconstituting either the liver or muscle clock alone (Fig. 3F). In contrast, and in accordance with our hypothesis, brain-RE mice had a notably improved glucose tolerance (Fig. 3F). This finding—that the central clock suffices for glucose homeostasis—complements recent work showing that it is necessary for glucose homeostasis (*28*). Together, these results underscore the importance of central processes in regulating glucose homeostasis and expands on the knowledge linking shift work to diabetes (*29*).

We next aimed to characterize the processes by which the central clock regulates systemic metabolism. We hypothesized that metabolic oscillations are driven by rhythmic feeding behavior; a process restored in the BRE mice (Extended Fig 2C). First, *ad hoc* analyses were performed on data from our published study (*22*) which assessed hepatic metabolite and transcript rhythms in KO mice that were night fed (NF), i.e. food was available only in the mouse’s active phase. An intact feeding/fasting cycle was sufficient for inducing ∼52% of oscillating liver metabolites, and 24% of oscillating liver transcripts, that are normally observed in WT mice (Fig. 4A), suggesting that the central clock and feed/fasting behavior impact on hepatic transcription to a similar degree. Strikingly, 56% of metabolic rhythms were rescued in *Bmal1*-KO mice simply by establishing a feeding rhythm for them (Table S3, Fig. 4B). Further, *ad libitum* fed brain-RE mice and NF-KO had almost an identical percent of recovery of the circadian metabolism and transcription of their respective WT littermates (Fig. 4C), showing that night feeding reinstated serum metabolite rhythms in KO mice to a level similar to that of brain-RE manipulation. The commonly oscillating metabolites between WT NF and KO NF were enriched for the acyl carnitine pathway (Extended Fig. 9A), in line with the shared pathways between brain-RE and WT mice. Nevertheless, the oscillating pattern of circulating metabolites in NF-KO and NF-WT mice were not identical to brain-RE or WT mice (fed *ad libitum*) (Fig. 4D): although we observed a 4-h delay in the early light-phase peak of the oscillating metabolites that were restored in brain-RE and KO NF, the uniquely rhythmic metabolites peaked in antiphase in the NF mice (Fig. 4E) and did not belong to the same pathways as in WT and brain-RE mice (Extended Data Fig. 9B, C). In addition, and in contrast to the higher amplitudes observed in brain-RE, the amplitude of the commonly oscillating metabolites was significantly lower in NF KO versus (*ad libitum* fed) WT mice (Fig. 4F). Finally, the same major metabolite classes displayed oscillations and peak times in NF-KO mice that were similar to those in WT and brain-RE mice, including a 4-h delay in the lipid peak in relation to WT (Fig. 4G). In sum, these data suggest that the central clock regulates a significant proportion of peripheral metabolic rhythms likely via feeding/fasting behavior, yet other clocks are required for a fully functioning circadian rhythmicity and for controlling certain aspects of the peripheral metabolic response to feeding, such as amplitude and phase.

**Figure 4.**
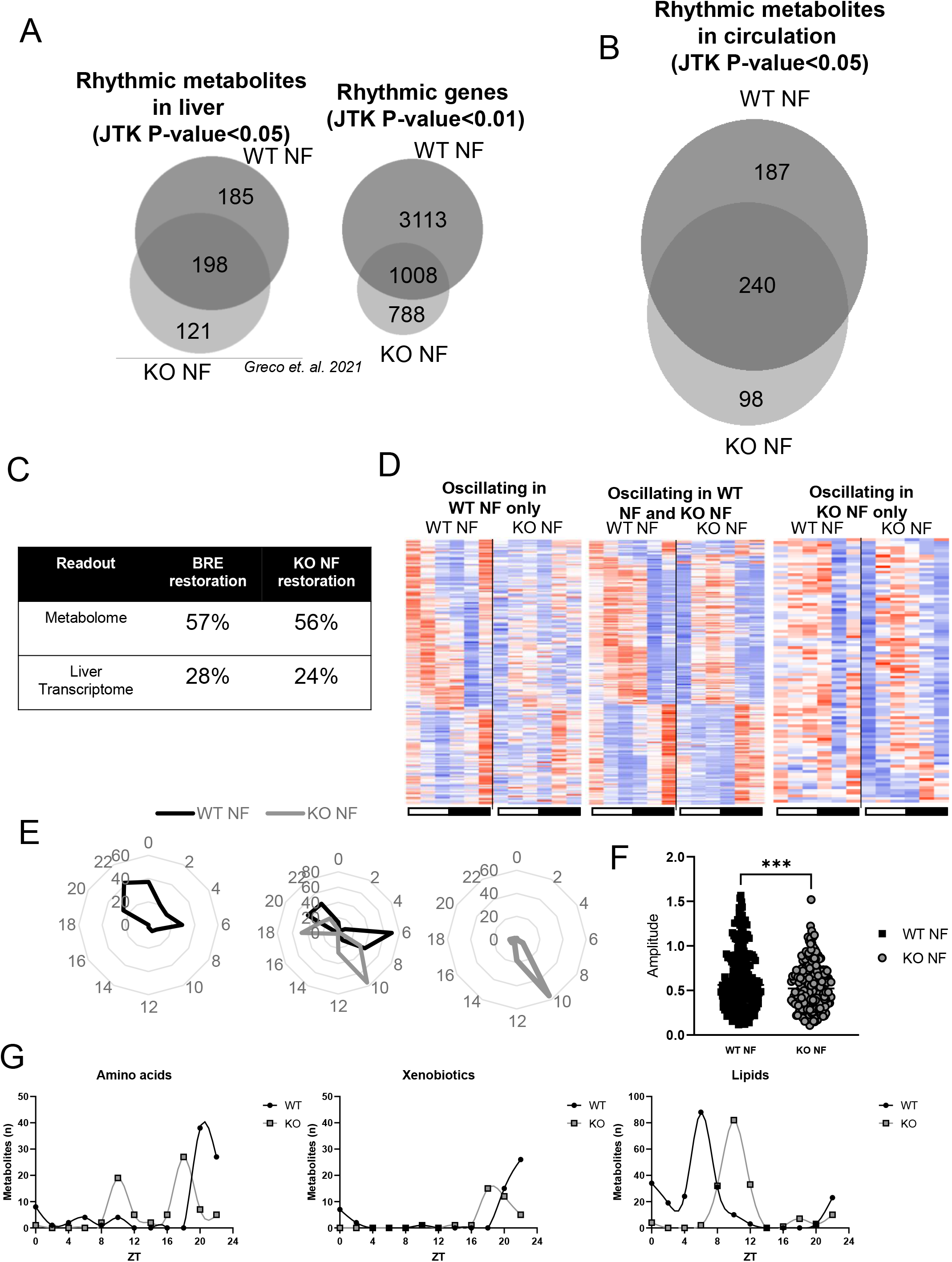

We are just starting to dissect the complex mechanisms governing circadian inter-organ communication (*4*). Here, we show that the central clock is sufficient for driving systemic rhythms to a large extent via regulation of feeding/fasting rhythms. It is known that food is a major synchronizing factor that contributes to the role of the SCN as a master pacemaker (*30*). Moreover, the notion that the SCN synchronizes some peripheral clocks via factors in circulation is supported by parabiosis experiments (*31*). However, interestingly, our data suggest that peripheral clocks are not only synchronized by food intake, but buffer and integrate the metabolic rhythms. Recently, we demonstrated an integrative role for the autonomous liver clock in the presence of feeding/fasting cycles (*22*), revealing cooperation between feeding/fasting cycles and the local clockwork in defining local rhythmic transcription. Supporting our findings, the hepatic clock has also been shown to shield untimely food intake and protect feeding-driven transcriptional rhythms in other peripheral tissues, such as the lung (*32*). It should be stressed that the phenotypes observed in brain-RE mice are independent of peripheral clocks; however, other neuronal clocks may explain some of the oscillations that are not restored (Extended Data Fig. 1C). In addition, it is possible that the *de novo* oscillating metabolites in the brain-RE and/or NF-KO mice drive aberrant rhythms in various tissues and thus require the liver clock to dampen their metabolic amplitude. In fact, this is supported by observations in muscles of the brain-RE mice, which display a substantial increase in oscillating transcripts (see accompanied manuscript Kumar et.al.). Taken together, this work presents the central clock as a key driver of systemic metabolic rhythms; future studies will elucidate how these metabolic signals are interpreted by various peripheral clocks, and how they can be best implemented into clinical therapies to treat morbidities associated to disrupted circadian rhythms (*33*).

## Acknowledgement

We thank all members of the Sassone-Corsi lab for helpful discussion and technical assistance. We greatly appreciate Melanie Oakes and Seung-Ah Chung at the UCI Genomics High-Throughput Facility. Furthermore, we thank the committee consisting of faculty in the Department of Biological Chemistry for their support after the passing of P.S.-C. P.P. was funded by The Wenner-Gren Foundations, The Foundation Blanceflor Boncompagni Ludovisi, née Bildt and Tore Nilsson Foundation for Medical Science. Funding for P.S.-C. was provided by the NIH (AG053592, DK114652), a Novo Nordisk Foundation Challenge Grant, and Institut National de la Sante et la Recherche Medicale (U1233 INSERM, France). Research in the S.A.B. lab is supported partially by the European Research Council (ERC) under the European Union’s Horizon 2020 research and innovation programme (Grant agreement No. 787041), the Government of Cataluña (SGR grant), the Government of Spain (MINECO), the La Marató/TV3 Foundation, the Foundation Lilliane Bettencourt, the Spanish Association for Cancer Research (AECC) and The Worldwide Cancer Research Foundation (WCRF). The IRB Barcelona is a Severo Ochoa Center of Excellence (MINECO award SEV-2015-0505). P.M.C acknowledges funding from MICINN-RTI2018-096068, ERC-2016-AdG-741966, LaCaixa-HEALTH-HR17- 00040, MDA, UPGRADE-H2020-825825, AFM, DPP-Spain, Fundació La MaratóTV3-80/19- 202021, MWRF, and María-de-Maeztu Program for Units of Excellence to UPF (MDM-2014- 0370) and the Severo- Ochoa Program for Centers of Excellence to CNIC (SEV-2015-0505). T.S. was supported by a Japan Society for the Promotion of Science (JSPS) fellowship. C.M.G. was funded by European Union’s Horizon 2020 research and innovation programme under the Marie Sklodowska-Curie grant agreement 749869. T.M. received funding from the European

Union’s Horizon 2020 research and innovation programme under the Marie Skłodowska-Curie grant agreement No. 754510. P.-S.W. is supported by grant RYC2019-026661-I funded by MCIN/AEI/10.13039/501100011033 and by “ESF Investing in your future”. We acknowledge predoctoral fellowships to M.C. from the National Institutes of Health (NIH) (GM117942), the American Heart Association (17PRE33410952), and the UCI School of Medicine Behrens Research Excellence Award. The work of S.C. and P.B. was in part supported by NIH grant NIH GM123558 to PB. We thank Veronica Raker for manuscript editing.

## Author contributions

P.P., J.G.S., K.B.K., P.M.-C., P. S.-C. and S.A.B. designed the study and analyzed the data. P.P., J.G.S., K.B.K., T.S., C.M.G., T.M., P.-S.W., V.Z., K.S. and M.C. performed experiments and/or interpreted the results. S.C. and P.B. performed the bioinformatics analyses. P.P. wrote the first version of the paper with feedback from all authors.

## Methods

### Animals

Mice were bred and housed in the animal facilities at the University of California–Irvine vivarium and Barcelona Science Park, Spain, in accordance with the guidelines of the Institutional Animal Care and Use Committee (IACUC) at UC-Irvine and European Union and Spanish regulations, respectively. Animal experiments were designed and conducted with consideration of the ARRIVE guidelines, the details of which were as follows. Identical experimental conditions were in place at both institutions, and all mice were derived from the same founder line. *Bmal1*-stop-FL mice were generated as described previously(*9, 10*). All experiments used 8–12-week-old male mice on a 12-h light/12-h dark cycle (or under constant darkness, for analyses of locomotor and indirect calorimetry) and fed *ad libitum*. Experimental mice were: knock-out (KO*) Bmal1*^stopFL/stopFL^, cre^−/−^; liver-RE (LRE) *Bmal1*^stopFL/stopFL^, *Alfp*-cre^−/tg^; muscle-RE (MRE) *Bmal1*^stopFL/stopFL^, *Hsa*-cre^−/tg^ and brain-RE (BRE) *Bmal1*^stopFL/stopFL^, *Syt10*- cre^−/tg^. Metabolomics analyses in *ad libitum* fed mice were performed using KO and wild-type (WT) littermates from all three genotypes: *Bmal1*^wt/wt^, *Alfp*-cre^−/tg^; *Bmal1*^wt/wt^, *Hsa*-cre^−/tg^ and *Bmal1*^wt/wt^, *Syt10*-cre^−/tg^. Metabolomics analyses in night-fed mice were performed in WT *Bmal1*^wt/wt^, *Alfp*-cre^−/tg^ and KO mice from the same colony. RNA-sequencing was performed using mice from the BRE colony, i.e., the *Bmal1*^wt/wt^, *Syt10*-cre^−/tg^ WTs. Mice were individually house for 2 weeks prior to sacrifice and tissue harvest.

### Global metabolomics analyses

Blood was collected from mice immediately after sacrifice and stored on ice for ∼30 min prior to centrifugation (3000 RPM for 10 min in 4 °C). Serum was collected in a fresh tube, snap frozen in liquid nitrogen and stored in –80 °C until shipment on dry ice to Metabolon Inc. Following receipt, samples were inventoried and immediately stored at –80°C. Each sample received was accessioned into the Metabolon LIMS system and was assigned by the LIMS a unique identifier that was associated with the original source identifier only. This identifier was used to track all sample handling, tasks, results, etc.

Samples were prepared using the automated MicroLab STAR® system from Hamilton Company. Several recovery standards were added prior to the first step in the extraction process for QC purposes. To remove protein, dissociate small molecules bound to protein or trapped in the precipitated protein matrix, and to recover chemically diverse metabolites, proteins were precipitated with methanol under vigorous shaking for 2 min (Glen Mills GenoGrinder 2000) followed by centrifugation. The resulting extract was divided into five fractions: two for analysis by two separate reverse phase (RP)/UPLC-MS/MS methods with positive ion mode electrospray ionization (ESI); one for analysis by RP/UPLC-MS/MS with negative ion mode ESI, one for analysis by HILIC/UPLC-MS/MS with negative ion mode ESI; and one reserved as a backup. Samples were placed briefly on a TurboVap® (Zymark) to remove the organic solvent. Sample extracts were stored overnight under nitrogen before preparation for analysis. Several types of controls were analysed in concert with the experimental samples: a pooled matrix sample generated by taking a small volume of each experimental sample (or alternatively, use of a pool of well-characterized human plasma), as a technical replicate throughout the data set; extracted water samples, as process blanks; and a cocktail of QC standards that were carefully chosen not to interfere with the measurement of endogenous compounds, which were spiked into every analyzed sample to allow instrument performance monitoring and to aid chromatographic alignment. Instrument variability was determined by calculating the median relative standard deviation (RSD) for the standards that were added to each sample prior to injection into the mass spectrometers. Overall process variability was determined by calculating the median RSD for all endogenous metabolites (i.e., non-instrument standards) present in 100% of the pooled matrix samples. Experimental samples were randomized across the platform run with QC samples spaced evenly among the injections.

All methods utilized a Waters ACQUITY ultra-performance liquid chromatography (UPLC) and a Thermo Scientific Q-Exactive high resolution/accurate mass spectrometer interfaced with a heated electrospray ionization (HESI-II) source and Orbitrap mass analyzer operated at 35,000 mass resolution. The sample extract was dried then reconstituted in solvents compatible to each of the four methods. Each reconstitution solvent contained a series of standards at fixed concentrations to ensure injection and chromatographic consistency. One aliquot was analyzed using acidic positive ion conditions, chromatographically optimized for more hydrophilic compounds. In this method, the extract was gradient eluted from a C18 column (Waters UPLC BEH C18-2.1×100 mm, 1.7 µm) using water and methanol, containing 0.05% perfluoropentanoic acid (PFPA) and 0.1% formic acid (FA). Another aliquot was also analyzed using acidic positive ion conditions; however, it was chromatographically optimized for more hydrophobic compounds. In this method, the extract was gradient eluted from the same afore mentioned C18 column using methanol, acetonitrile, water, 0.05% PFPA and 0.01% FA and was operated at an overall higher organic content. Another aliquot was analyzed using basic negative ion optimized conditions using a separate dedicated C18 column. The basic extracts were gradient eluted from the column using methanol and water but using 6.5 mM ammonium bicarbonate, pH 8. The fourth aliquot was analysed via negative ionization following elution from a HILIC column (Waters UPLC BEH Amide 2.1×150 mm, 1.7 µm) using a gradient consisting of water and acetonitrile with 10 mM ammonium formate, pH 10.8. The MS analysis alternated between MS and data-dependent MSn scans using dynamic exclusion. The scan range varied slighted between methods but covered 70-1000 m/z.

Raw data was extracted, peak-identified and QC processed using Metabolon’s hardware and software. These systems are built on a web-service platform utilizing Microsoft’s .NET technologies, which run on high-performance application servers and fiber-channel storage arrays in clusters to provide active failover and load-balancing. Compounds were identified by comparison to library entries of purified standards or recurrent unknown entities. Metabolon maintains a library based on authenticated standards that contains the retention time/index (RI), mass to charge ratio (m/z), and chromatographic data (including MS/MS spectral data) on all molecules present in the library. Furthermore, biochemical identifications are based on three criteria: retention index within a narrow RI window of the proposed identification, accurate mass match to the library +/– 10 ppm, and the MS/MS forward and reverse scores between the experimental data and authentic standards. The MS/MS scores are based on a comparison of the ions present in the experimental spectrum to the ions present in the library spectrum. While there may be similarities between these molecules based on one of these factors, the use of all three data points can be utilized to distinguish and differentiate biochemicals. More than 3300 commercially available, purified standard compounds have been acquired and registered into LIMS for analysis on all platforms for determination of their analytical characteristics. A variety of curation procedures was carried out to ensure that a high-quality dataset was available for statistical analysis and data interpretation. The QC and curation processes were designed to ensure accurate and consistent identification of true chemical entities, and to remove those representing system artifacts, mis-assignments, and background noise. Metabolon data analysts use proprietary visualization and interpretation software to confirm the consistency of peak identification among the various samples. Library matches for each compound were checked for each sample and corrected if necessary.

Peaks were quantified using area-under-the-curve. A data normalization step was performed to correct variation resulting from instrument inter-day tuning differences. Essentially, each compound was corrected in run-day blocks by registering the medians to equal one and normalizing each data point proportionately. Furthermore, the data were normalized to the input volume of serum. Finally, the results were analyzed and clear outliers were excluded. Metabolic circadian rhythms were assessed using JTK-CYCLE(*11*) and *P*-values < 0.05 were considered significant.

### Indirect calorimetry and food intake

Indirect calorimetry was performed with negative flow Oxymax-CLAMS (Columbus Instruments, Columbus, OH) hardware system cages. Mice were given a 24-h acclimation period to the metabolic cage. Measurements of energy expenditure, oxygen respiration (VO_2_), carbon dioxide respiration (VCO_2_) and food intake were taken every 10 min for 2 consecutive days after the 24-h acclimation period at room temperature. The respiratory exchange ratio (RER = VCO2/VO2) was calculated by the accompanying Oxymax software. The measurements were performed under 12-h light/dark cycles or in constant darkness.

### Locomotor activity

Locomotor activity of individually housed mice was measured using optical beam motion detection (Starr Life Sciences). Data were collected using Minimitter VitalView 5.0 data acquisition software and analysed using Clocklab (Actimetrics). Mouse were housed under 12-h light/dark cycles for the initial 2 weeks, followed by a transition to 3 weeks in constant darkness.

### Oral glucose tolerance test

Body weights were measured, and 2 g kg^−1^ glucose was delivered orally. Blood glucose measurements were taken pre-injection and at 15, 30, 60 and 120 min post-glucose injection at ZT14 using a Accu-Chek Aviva glucose meter kit.

### Protein extraction and Western blot

Frozen tissue pieces were placed in 1.5 ml Eppendorf tubes with RIPA buffer (50 mM Tris-HCl [pH 8.0], 150 mM NaCl, 1% NP-40, 0.5% sodium deoxycholate, 0.1% SDS, 5 mM MgCl_2_ and 1 mM PMSF) supplemented with Protease Inhibitor Cocktail (Roche), 1 mM DTT, 20 mM NaF, 10 mM nicotinamide and 330 nM trichostatin A (T8552, Sigma). Samples were homogenized using a VWR 200 homogenizer followed by a 10-sec sonication at 60%. Tissue lysates were centrifugated at 13200 rpm for 15 min at 4 °C. Supernatants were transferred to a new tube and an aliquot were used for protein quantification using the Bradford assay (Bio-Rad). A total of 10-20 μg of protein lysates were separated on 8% gels by SDS-PAGE and transferred to nitrocellulose membranes. Membranes were incubated with primary anti-BMAL1, Abcam, ab93806 and anti-P84, Genetex, GTX70220 antibodies overnight at 4 °C, and with peroxidase- conjugated secondary antibodies (Anti-mouse IgG, HRP conjugate, EMD Millipore, AP160P; Anti-rabbit IgG, HRP-linked, EMD Millipore, 12-348) for 1 h at room temperature. Protein bands were visualized by chemiluminescent HRP substrate (WBKLS0500, EMD Millipore). Blots were developed with autoradiography films (HyBlot CL, Denville Scientific) or ChemiDoc™ MP Imaging System (Bio-Rad Laboratories).

### Immunohistochemistry

Mouse brains were frozen at –80 °C isopentane and sectioned in 10µm thick sections using Leica CM1950 Cryostat. Sections were fixed with ice cold 4% PFA for 20 min and washed three times with 1× PBS. Samples were then permeabilized (1× PBS, 0.3% Triton X-100) for 15 min at room temperature (RT) and blocked (in 1× PBS, 5% BSA and 10% normal goat serum) for 2 h at RT, and then incubated with a BMAL1 antibody (Novus, NB100-2288) diluted in blocking buffer overnight at 4°C. Samples were washed three times with 1× PBS and incubated with Alexa Fluor 488 goat anti-rabbit IgG (Invitrogen, A-11008) for 2 h at RT. After three washes with 1× PBS, nuclei were stained using DRAQ7 (Biostatus) for 15 min at RT and subsequently washed twice with 1× PBS.

### RNA extraction and RNA-sequencing and analysis

Total RNA was extracted from liver with TRIzol (Invitrogen) and then precipitated with isopropanol and ethanol. Total RNA was monitored for quality control using the Agilent Bioanalyzer Nano RNA chip and Nanodrop absorbance ratios for 260/280 nm and 260/230 nm.

Libraries were constructed according to the Illumina TruSeq® Stranded mRNA Sample Preparation Guide. The input quantity for total RNA was 1000 ng, and mRNA was enriched using oligo dT magnetic beads. The enriched mRNA was chemically fragmented for 3 min. First-strand synthesis used random primers and reverse transcriptase to make cDNA. After second-strand synthesis, the (ds)-cDNA was cleaned using AMPure XP beads, cDNA was end- repaired and the 3′ ends were adenylated. Illumina barcoded adapters were ligated on the ends, and the adapter-ligated fragments were enriched by nine cycles of PCR. The resulting libraries were validated by qPCR and sized using a Agilent Bioanalyzer DNA high-sensitivity chip. The concentrations for the libraries were normalized and then multiplexed together. The multiplexed libraries were sequenced on paired end 100 cycles chemistry on the Novaseq 6000. The Novaseq control software NVCS ver 1.7.0 was used with real-time analysis software, RTA 3.4.4.

Reads from each replicate experiment were aligned to the reference genome assembly mm10 and to the corresponding transcriptome using TopHat. Subsequently, gene expression levels were computed from the read alignment results using Cufflinks, another tool in the Tuxedo protocol. This protocol outputs the FPKM values for each gene of each replicate. To determine periodicity of genes, the JTK-CYCLE(*11*) algorithm was used. The output of this algorithm includes the amplitude, phase and p-value for each transcript. Genes were considered circadian if their *P*- value output by JTK-CYCLE was < 0.01. Heatmaps of circadian transcripts were generated using the R package gplots v3.0.3, in which the rows are sorted by the JTK_CYCLE output phase and row z-score normalized. Gene ontology and pathway analysis was performed using ToppFun (https://toppgene.cchmc.org/).

### Statistics

For each experiment, the number of biological replicates, statistical test, significance threshold and visual representation information (i.e., mean, SEM, etc. of graphs) are given in the Fig. legends.

### Data availability

The metabolomics and transcriptomics data and its analyses is available from the CircadiOmics web portal (*34, 35*).

**Figure S1.**
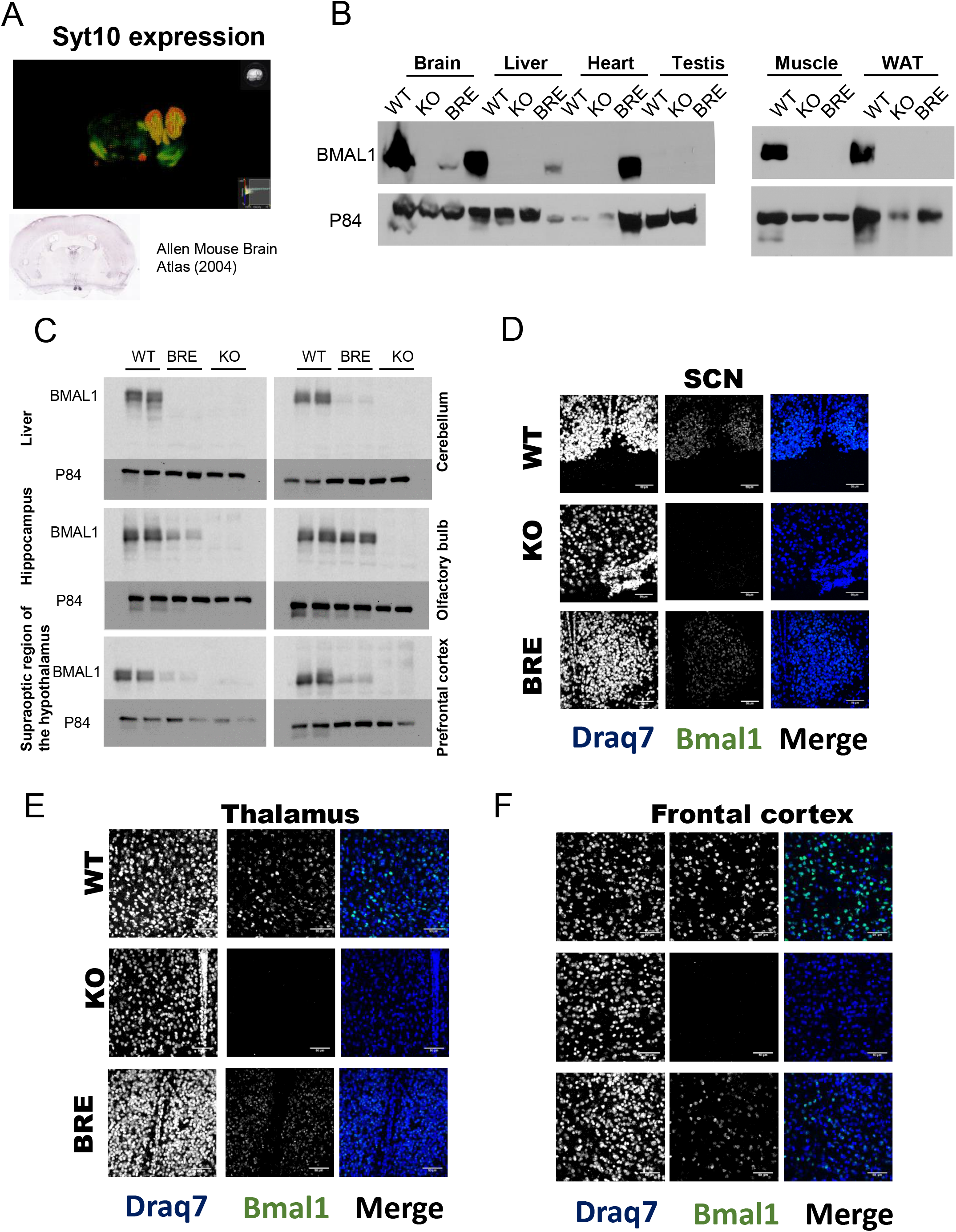

**Figure S2.**
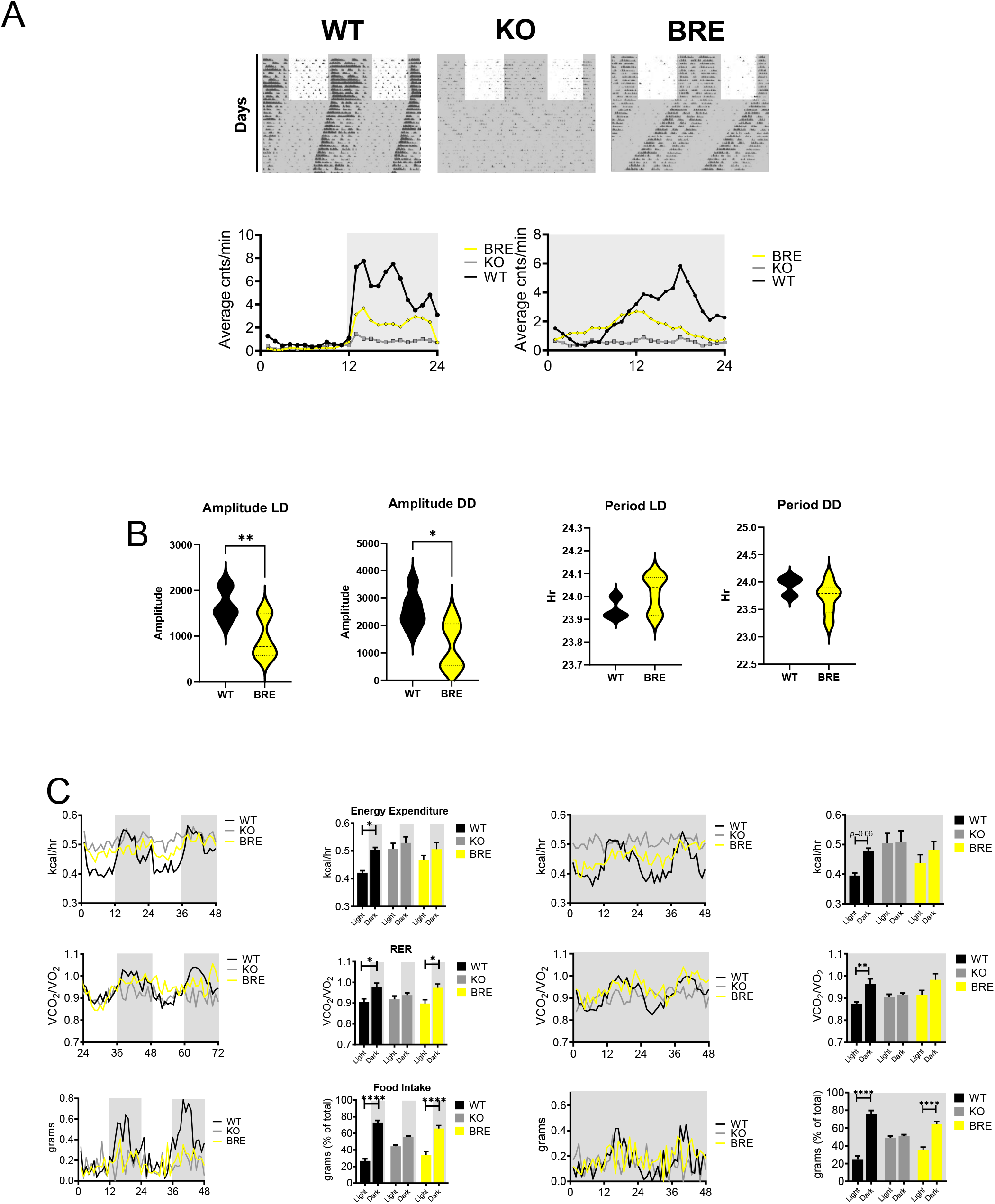

**Figure S3.**
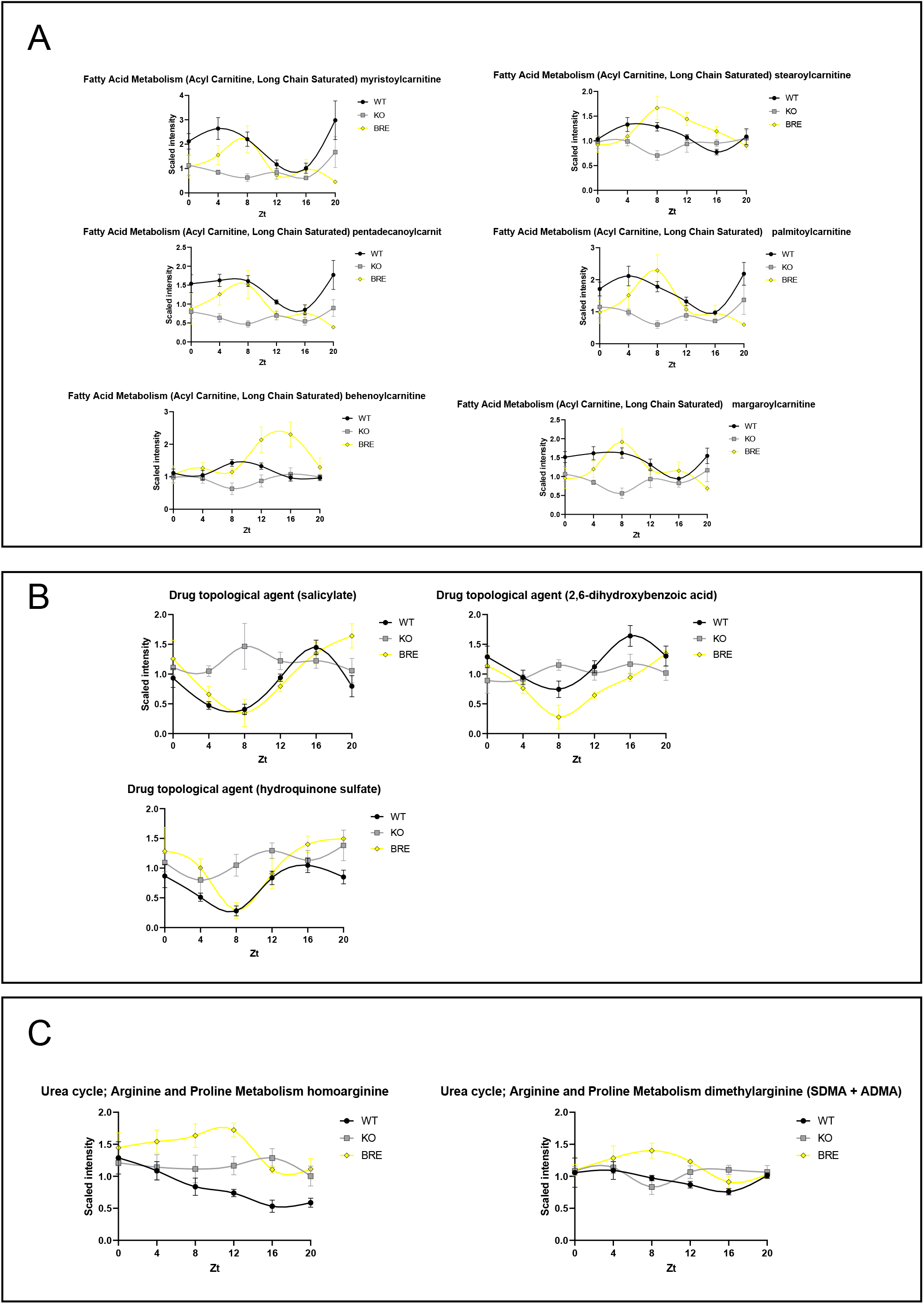

**Figure S4.**
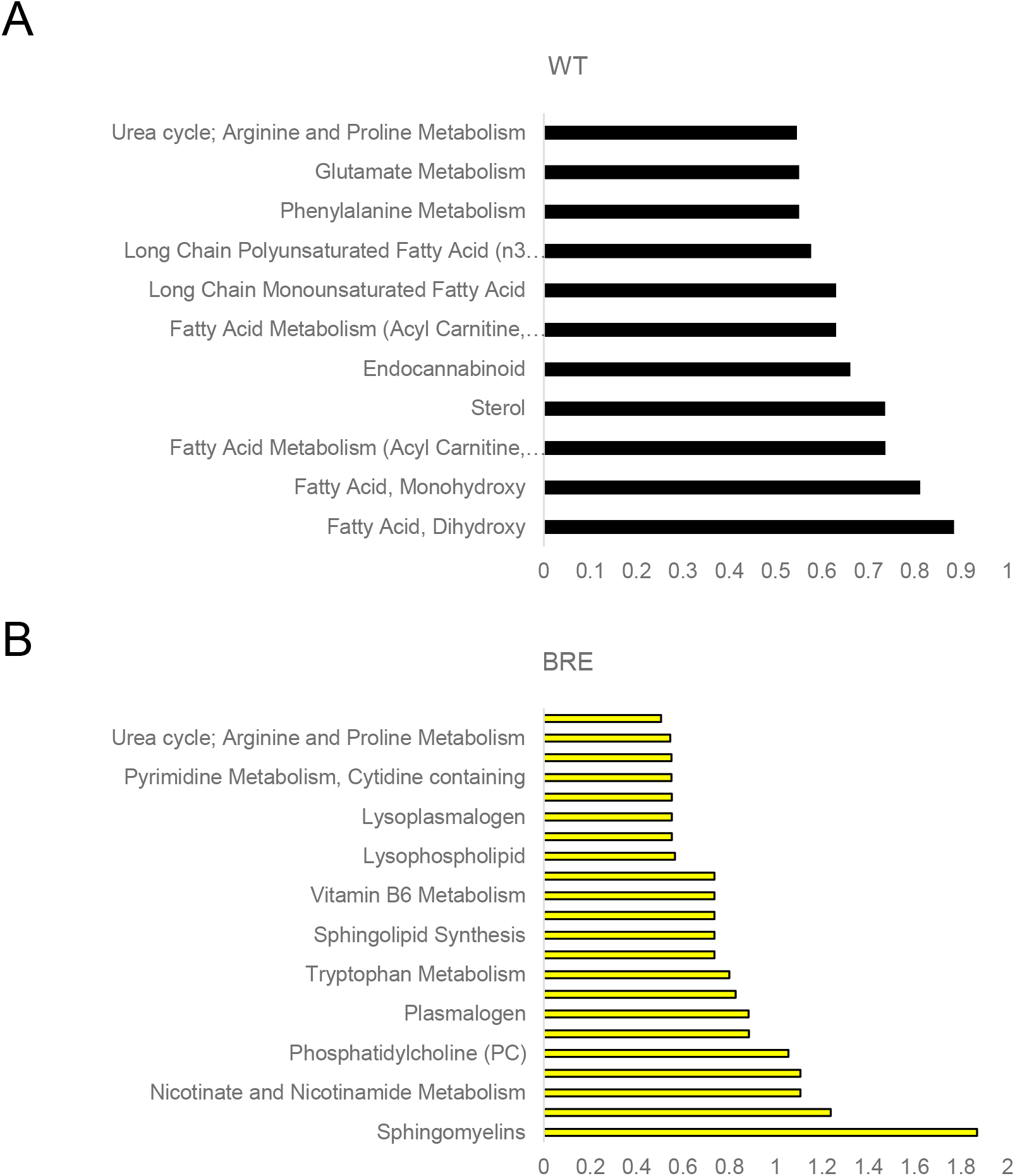

**Figure S5.**
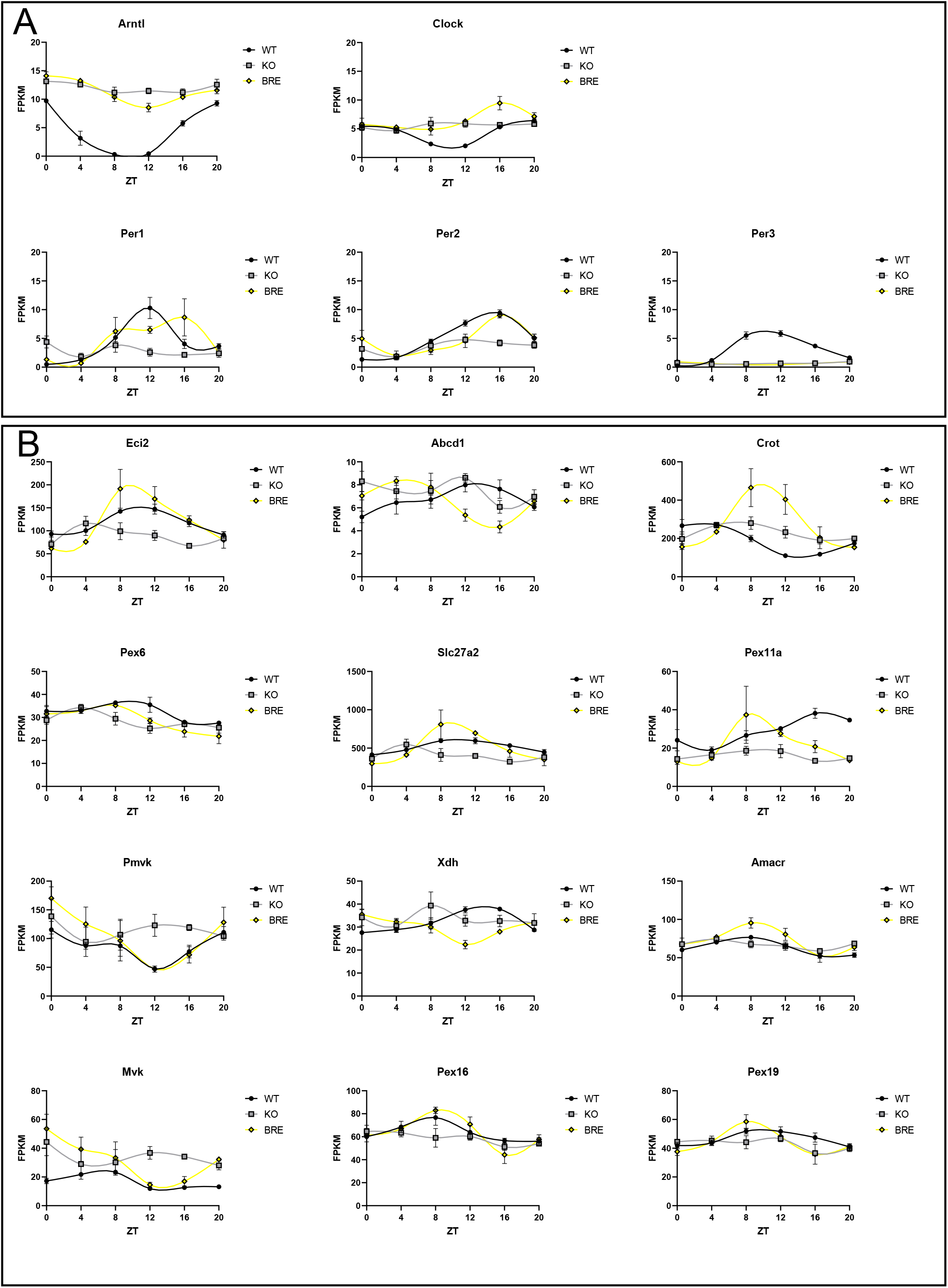

**Figure S6.**
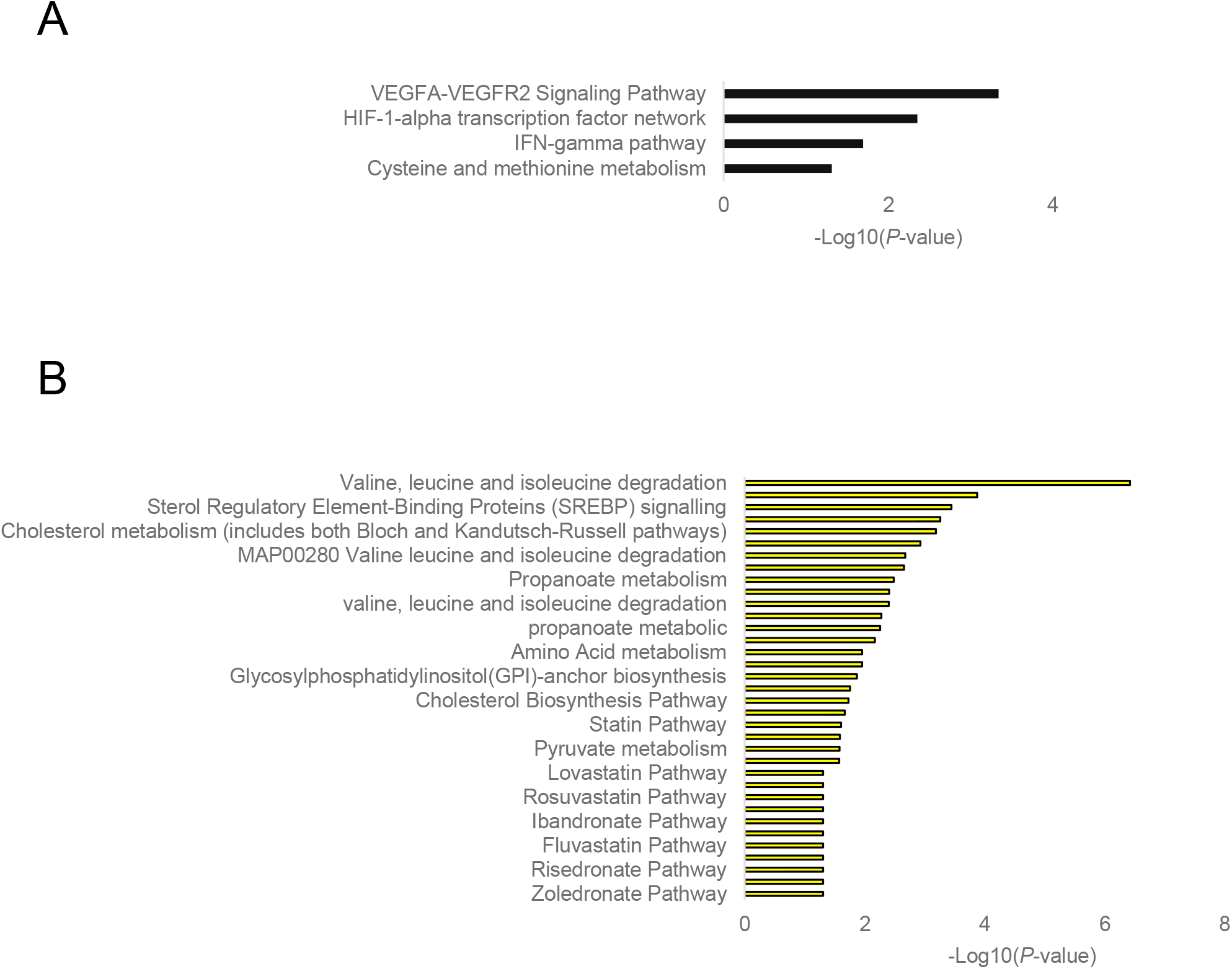

**Figure S7.**
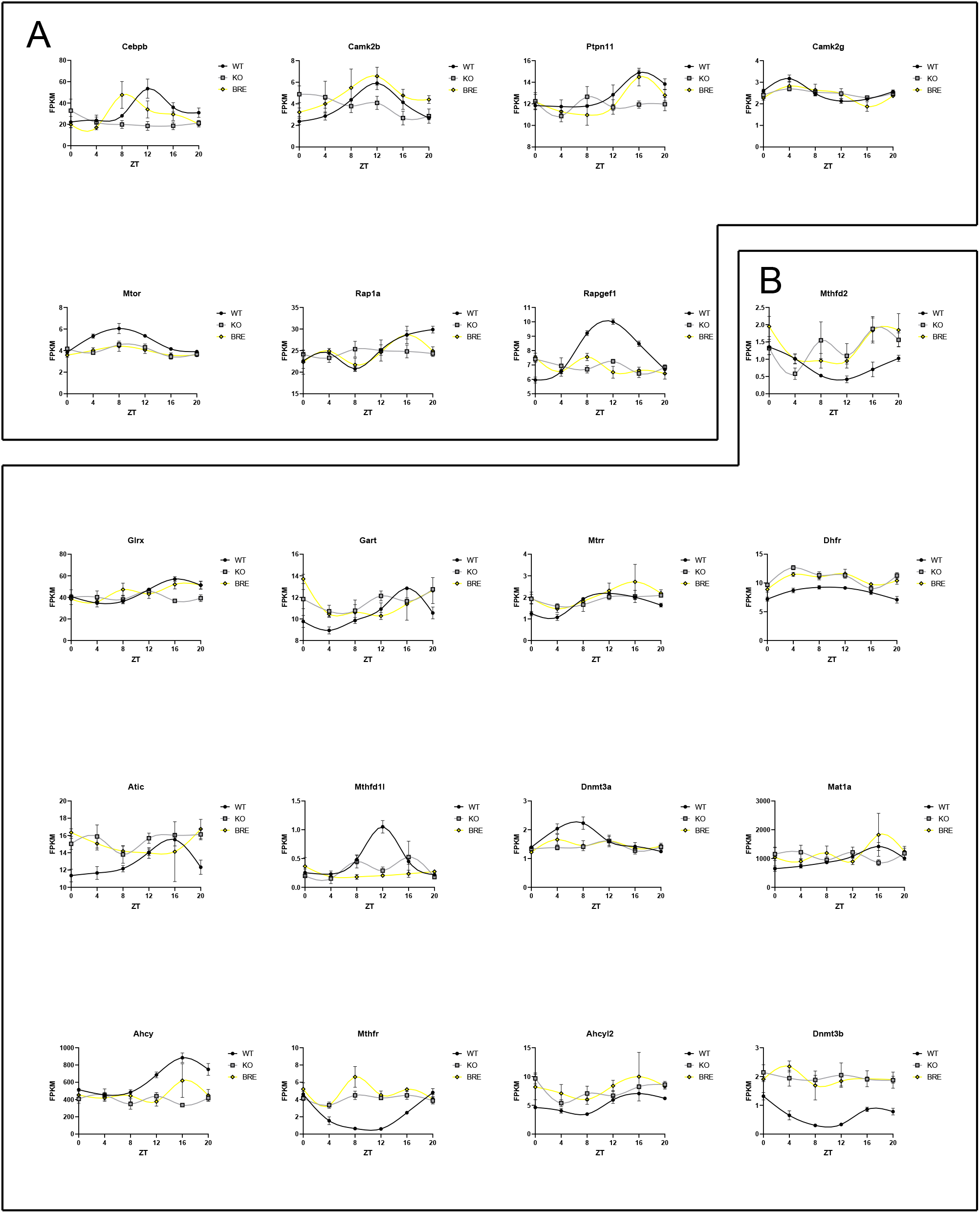

**Figure S8.**
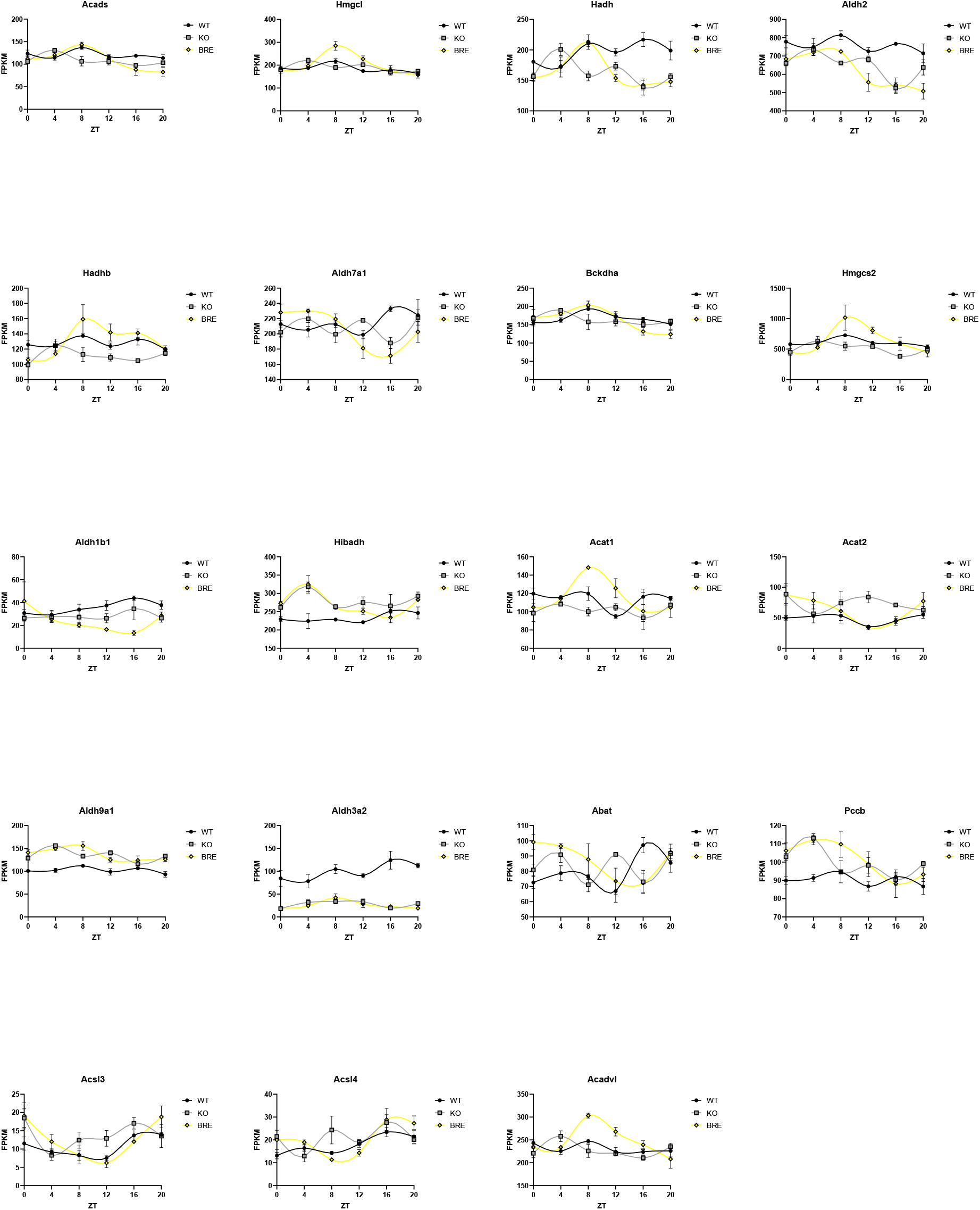

**Figure S9.**
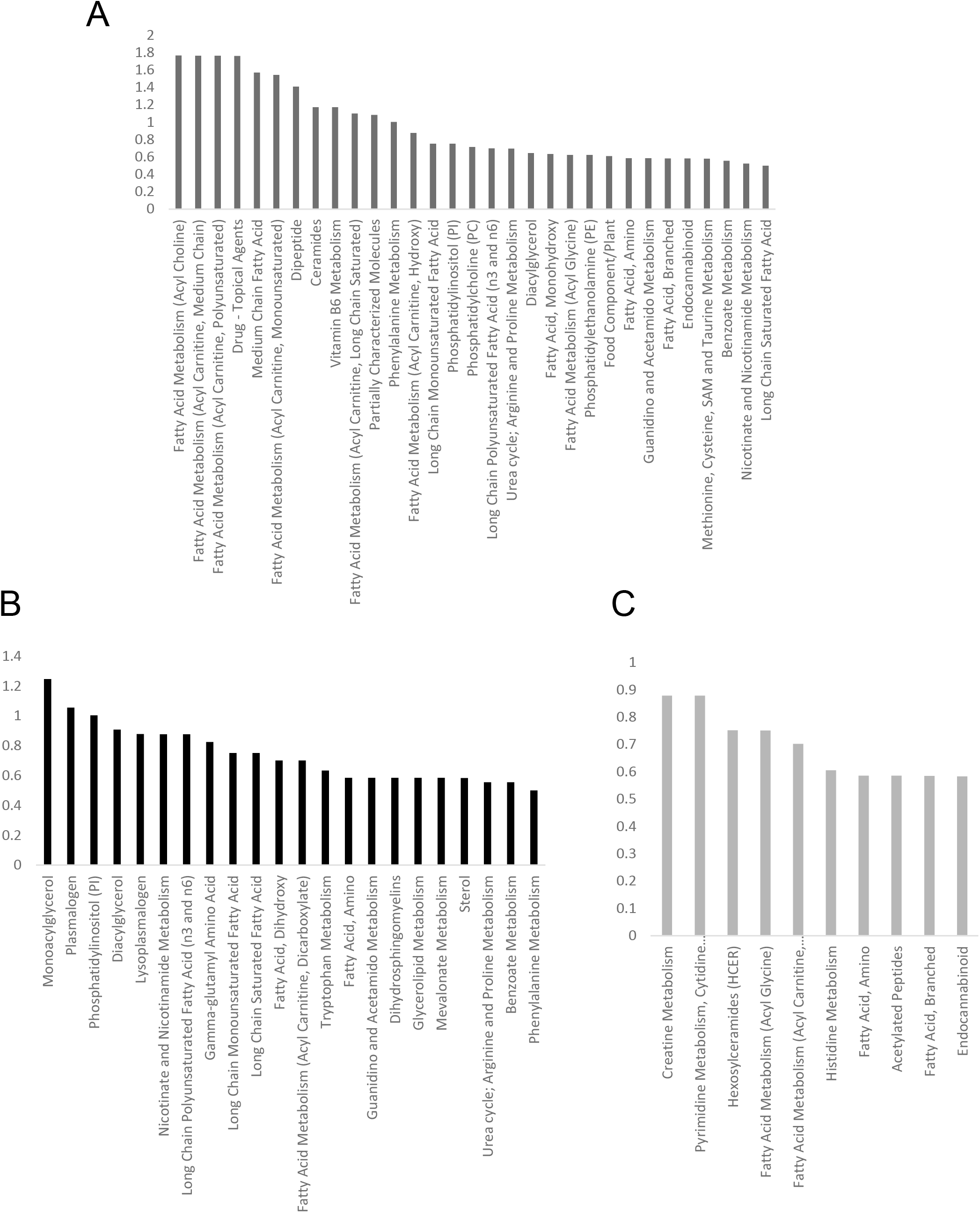

## References

1. J. S. Takahashi, Transcriptional architecture of the mammalian circadian clock. Nat. Rev. Genet. 18 (2017), pp. 164–179.

2. K. B. Koronowski, P. Sassone-Corsi, Communicating clocks shape circadian homeostasis. Science (80-.). 371, eabd0951 (2021).

3. J. Ye, R. Medzhitov, Control strategies in systemic metabolism. Nat. Metab. 1 (2019), pp. 947–957.

4. H. Reinke, G. Asher, Crosstalk between metabolism and circadian clocks. Nat. Rev. Mol. Cell Biol. 20 (2019), pp. 227–241.

5. K. A. Dyar, D. Lutter, A. Artati, N. J. Ceglia, Y. Liu, D. Armenta, M. Jastroch, S. Schneider, S. de Mateo, M. Cervantes, S. Abbondante, P. Tognini, R. Orozco-Solis, K. Kinouchi, C. Wang, R. Swerdloff, S. Nadeef, S. Masri, P. Magistretti, V. Orlando, E. Borrelli, N. H. Uhlenhaut, P. Baldi, J. Adamski, M. H. Tschöp, K. Eckel-Mahan, P. Sassone-Corsi, Atlas of Circadian Metabolism Reveals System-wide Coordination and Communication between Clocks. Cell. 174, 1571–1585.e11 (2018).

6. P. L. Lowrey, J. S. Takahashi, in Advances in Genetics (Academic Press Inc., 2011), vol. 74, pp. 175–230.

7. J. Bass, J. S. Takahashi, Circadian integration of metabolism and energetics. Science (80-.). 330 (2010), pp. 1349–1354.

8. S. Panda, Circadian physiology of metabolism. Science. 354, 1008–1015 (2016).

9. K. B. Koronowski, K. Kinouchi, P.-S. Welz, J. G. Smith, V. M. Zinna, J. Shi, M. Samad, S. Chen, C. N. Magnan, J. M. Kinchen, W. Li, P. Baldi, S. A. Benitah, P. Sassone-Corsi, Defining the Independence of the Liver Circadian Clock. Cell. 177, 1448–1462.e14 (2019).

10. P.-S. Welz, V. M. Zinna, A. Symeonidi, K. B. Koronowski, K. Kinouchi, J. G. Smith, I. M. Guillén, A. Castellanos, S. Furrow, F. Aragón, G. Crainiciuc, N. Prats, J. M. Caballero, A. Hidalgo, P. Sassone-Corsi, S. A. Benitah, BMAL1-Driven Tissue Clocks Respond Independently to Light to Maintain Homeostasis. Cell. 177, 1436–1447.e12 (2019).

11. M. E. Hughes, J. B. Hogenesch, K. Kornacker, JTK-CYCLE: An efficient nonparametric algorithm for detecting rhythmic components in genome-scale data sets. J. Biol. Rhythms. 25, 372–380 (2010).

12. Y. Nakahata, S. Sahar, G. Astarita, M. Kaluzova, P. Sassone-Corsi, Circadian Control of the NAD+ Salvage Pathway by CLOCK-SIRT1. Science (80-.). 324, 654–657 (2009).

13. Y. Nakahata, M. Kaluzova, B. Grimaldi, S. Sahar, J. Hirayama, D. Chen, L. P. Guarente, P. Sassone-Corsi, The NAD+-Dependent Deacetylase SIRT1 Modulates CLOCK-Mediated Chromatin Remodeling and Circadian Control. Cell. 134, 329–340 (2008).

14. H. Abe, S. Honma, H. Ohtsu, K. I. Honma, Circadian rhythms in behavior and clock gene expressions in the brain of mice lacking histidine decarboxylase. Mol. Brain Res. 124, 178–187 (2004).

15. K. A. Dyar, M. J. Hubert, A. A. Mir, S. Ciciliot, D. Lutter, F. Greulich, F. Quagliarini, M. Kleinert, K. Fischer, T. O. Eichmann, L. E. Wright, M. I. Peña Paz, A. Casarin, V. Pertegato, V. Romanello, M. Albiero, S. Mazzucco, R. Rizzuto, L. Salviati, G. Biolo, B. Blaauw, S. Schiaffino, N. H. Uhlenhaut, Transcriptional programming of lipid and amino acid metabolism by the skeletal muscle circadian clock. PLoS Biol. 16, e2005886 (2018).

16. D. R. Weaver, The suprachiasmatic nucleus: a 25-year retrospective. J. Biol. Rhythms. 13, 100–12 (1998).

17. E. S. Lein, M. J. Hawrylycz, N. Ao, M. Ayres, A. Bensinger, A. Bernard, A. F. Boe, M. S. Boguski, K. S. Brockway, E. J. Byrnes, L. Chen, L. Chen, T. M. Chen, M. C. Chin, J. Chong, B. E. Crook, A. Czaplinska, C. N. Dang, S. Datta, N. R. Dee, A. L. Desaki, T. Desta, E. Diep, T. A. Dolbeare, M. J. Donelan, H. W. Dong, J. G. Dougherty, B. J. Duncan, A. J. Ebbert, G. Eichele, L. K. Estin, C. Faber, B. A. Facer, R. Fields, S. R. Fischer, T. P. Fliss, C. Frensley, S. N. Gates, K. J. Glattfelder, K. R. Halverson, M. R. Hart, J. G. Hohmann, M. P. Howell, D. P. Jeung, R. A. Johnson, P. T. Karr, R. Kawal, J. M. Kidney, R. H. Knapik, C. L. Kuan, J. H. Lake, A. R. Laramee, K. D. Larsen, C. Lau, T. A. Lemon, A. J. Liang, Y. Liu, L. T. Luong, J. Michaels, J. J. Morgan, R. J. Morgan, M. T. Mortrud, N. F. Mosqueda, L. L. Ng, R. Ng, G. J. Orta, C. C. Overly, T. H. Pak, S. E. Parry, S. D. Pathak, O. C. Pearson, R. B. Puchalski, Z. L. Riley, H. R. Rockett, S. A. Rowland, J. J. Royall, M. J. Ruiz, N. R. Sarno, K. Schaffnit, N. V. Shapovalova, T. Sivisay, C. R. Slaughterbeck, S. C. Smith, K. A. Smith, B. I. Smith, A. J. Sodt, N. N. Stewart, K. R. Stumpf, S. M. Sunkin, M. Sutram, A. Tam, C. D. Teemer, C. Thaller, C. L. Thompson, L. R. Varnam, A. Visel, R. M. Whitlock, P. E. Wohnoutka, C. K. Wolkey, V. Y. Wong, M. Wood, M. B. Yaylaoglu, R. C. Young, B. L. Youngstrom, X. F. Yuan, B. Zhang, T. A. Zwingman, A. R. Jones, Genome-wide atlas of gene expression in the adult mouse brain. Nature. 445, 168–176 (2007).

18. J. Husse, X. Zhou, A. Shostak, H. Oster, G. Eichele, Synaptotagmin10-Cre, a Driver to Disrupt Clock Genes in the SCN. J. Biol. Rhythms. 26, 379–389 (2011).

19. K. A. Dyar, S. Ciciliot, L. E. Wright, R. S. Biensø, G. M. Tagliazucchi, V. R. Patel, M. Forcato, M. I. P. Paz, A. Gudiksen, F. Solagna, M. Albiero, I. Moretti, K. L. Eckel-Mahan, P. Baldi, P. Sassone-Corsi, R. Rizzuto, S. Bicciato, H. Pilegaard, B. Blaauw, S. Schiaffino, Muscle insulin sensitivity and glucose metabolism are controlled by the intrinsic muscle clock. Mol. Metab. 3, 29–41 (2014).

20. K. A. Lamia, K. F. Storch, C. J. Weitz, Physiological significance of a peripheral tissue circadian clock. Proc. Natl. Acad. Sci. U. S. A. 105, 15172–15177 (2008).

21. R. Lin, Y. Mo, H. Zha, Y. Xu, Y. Xiong, K.-L. Guan Correspondence, CLOCK Acetylates ASS1 to Drive Circadian Rhythm of Ureagenesis. Mol. Cell. 68 (2017), doi:10.1016/j.molcel.2017.09.008.

22. C. M. Greco, K. B. Koronowski, J. G. Smith, J. Shi, P. Kunderfranco, R. Carriero, S. Chen, M. Samad, P.-S. Welz, V. M. Zinna, T. Mortimer, S. K. Chun, K. Shimaji, T. Sato, P. Petrus, A. Kumar, M. Vaca-Dempere, O. Deryagian, C. Van, J. M. M. Kuhn, D. Lutter, M. M. Seldin, S. Masri, W. Li, P. Baldi, K. A. Dyar, P. Muñoz-Cánoves, S. A. Benitah, P. Sassone-Corsi, Integration of feeding behavior by the liver circadian clock reveals network dependency of metabolic rhythms. Sci. Adv. 7 (2021), doi:10.1126/SCIADV.ABI7828.

23. B. Kornmann, O. Schaad, H. Bujard, J. S. Takahashi, U. Schibler, System-driven and oscillator-dependent circadian transcription in mice with a conditionally active liver clock. PLoS Biol. 5, 0179–0189 (2007).

24. C. B. Peek, D. C. Levine, J. Cedernaes, A. Taguchi, Y. Kobayashi, S. J. Tsai, N. A. Bonar, M. R. McNulty, K. M. Ramsey, J. Bass, Circadian Clock Interaction with HIF1α Mediates Oxygenic Metabolism and Anaerobic Glycolysis in Skeletal Muscle. Cell Metab. 25, 86–92 (2017).

25. R. G. Carroll, G. A. Timmons, M. P. Cervantes-Silva, O. D. Kennedy, A. M. Curtis, Immunometabolism around the Clock. Trends Mol. Med. 25 (2019), pp. 612–625.

26. C. M. Greco, M. Cervantes, J.-M. Fustin, K. Ito, N. Ceglia, M. Samad, J. Shi, K. B. Koronowski, I. Forne, S. Ranjit, J. Gaucher, K. Kinouchi, R. Kojima, E. Gratton, W. Li, P. Baldi, A. Imhof, H. Okamura, P. Sassone-Corsi, S-adenosyl-l-homocysteine hydrolase links methionine metabolism to the circadian clock and chromatin remodeling. Sci. Adv. 6, eabc5629 (2020).

27. M. G. Schooneman, F. M. Vaz, S. M. Houten, M. R. Soeters, Acylcarnitines: Reflecting or inflicting insulin resistance? Diabetes. 62 (2013), pp. 1–8.

28. G. Ding, X. Li, X. Hou, W. Zhou, Y. Gong, F. Liu, Y. He, J. Song, J. Wang, P. Basil, W. Li, S. Qian, P. Saha, J. Wang, C. Cui, T. Yang, K. Zou, Y. Han, C. I. Amos, Y. Xu, L. Chen, Z. Sun, REV-ERB in GABAergic neurons controls diurnal hepatic insulin sensitivity. Nature. 592, 763–767 (2021).

29. J. Qian, F. A. J. L. Scheer, Circadian System and Glucose Metabolism: Implications for Physiology and Disease. Trends Endocrinol. Metab. 27 (2016), pp. 282–293.

30. G. Asher, P. Sassone-Corsi, Time for food: the intimate interplay between nutrition, metabolism, and the circadian clock. Cell. 161, 84–92 (2015).

31. H. Guo, J. M. K. Brewer, A. Champhekar, R. B. S. Harris, E. L. Bittman, Differential control of peripheral circadian rhythms by suprachiasmatic-dependent neural signals. Proc. Natl. Acad. Sci. U. S. A. 102, 3111–3116 (2005).

32. G. Manella, E. Sabath, R. Aviram, V. Dandavate, S. Ezagouri, M. Golik, Y. Adamovich, G. Asher, The liver-clock coordinates rhythmicity of peripheral tissues in response to feeding. Nat. Metab., 1–14 (2021).

33. C. R. Cederroth, U. Albrecht, J. Bass, S. A. Brown, J. Dyhrfjeld-Johnsen, F. Gachon, C. B. Green, M. H. Hastings, C. Helfrich-Förster, J. B. Hogenesch, F. Lévi, A. Loudon, G. B. Lundkvist, J. H. Meijer, M. Rosbash, J. S. Takahashi, M. Young, B. Canlon, Medicine in the Fourth Dimension. Cell Metab. 30 (2019), pp. 238–250.

34. N. Ceglia, Y. Liu, S. Chen, F. Agostinelli, K. Eckel-Mahan, P. Sassone-Corsi, P. Baldi, CircadiOmics: circadian omic web portal. Nucleic Acids Res. 46, W157–W162 (2018).

35. V. R. Patel, K. Eckel-Mahan, P. Sassone-Corsi, P. Baldi, CircadiOmics: integrating circadian genomics, transcriptomics, proteomics and metabolomics. Nat. Methods 2012 98. 9, 772–773 (2012).

